# Structural basis of human replisome progression into a nucleosome

**DOI:** 10.1101/2025.04.04.647053

**Authors:** Felix Steinruecke, Jonathan W. Markert, Lucas Farnung

## Abstract

Epigenetic inheritance requires the transfer of parental histones to newly synthesized DNA during eukaryotic chromosome replication, yet the structural mechanisms underlying replisome engagement with nucleosomes remain unclear. Here we establish an *in vitro* chromatin replication system and report four cryo-EM structures of the human replisome in complex with a parental nucleosome. The structures capture distinct states of nucleosomal DNA unwrapping and nucleosome integrity during nucleosome disassembly by the encroaching replisome.

## Main

In eukaryotes, DNA replication occurs in the context of chromatin^1,2^, the fundamental unit of which is the nucleosome core particle^3,4^. Parental nucleosomes, composed of two copies of histone proteins H2A, H2B, H3, and H4 wrapped by 145-147 base pairs (bp) of DNA, must be disrupted to enable DNA unwinding and daughter strand synthesis by the replisome^5^. To ensure the inheritance of epigenetic information, parental histones must be transferred to and reassembled into nucleosomes on the newly synthesized leading and lagging DNA strands^6^. However, the mechanism by which the replisome, comprising the CDC45-MCM-GINS (CMG) helicase, DNA polymerases, and accessory factors^7^, engages and disrupts parental nucleosomes to facilitate their transfer remains unresolved^8^. Here, we established a reconstituted chromatin replication system and determined cryo-electron microscopy (cryo-EM) structures of the human replisome during its initial engagement with and progression into a parental nucleosome. Our findings reveal a stepwise mechanism of nucleosomal DNA unwrapping from the histone core at the front of the replisome.

## Results and discussion

To investigate how the replisome traverses nucleosomes, we reconstituted CMG-dependent leading-strand DNA replication on a nucleosomal substrate *in vitro*. We recombinantly purified components of the human replication machinery^9^ (Extended Data Fig. 1a) and prepared a forked DNA substrate consisting of 125 nucleotide (nt) leading and 20 nt lagging single-stranded DNA arms and a 145 bp Widom 601 nucleosome positioning sequence, separated by an additional 71 bp of double-stranded DNA (Extended Data Fig. 1b,c). A nucleosome was assembled on the Widom 601 sequence. Helicase-competent CMG (Extended Data Fig. 1d,e) was loaded onto the forked substrate and a fluorescently labeled DNA primer was annealed to the leading strand template (Extended Data Fig. 1f). TIMELESS-TIPIN, AND-1, CLASPIN, DNA polymerase ε (Pol ε), RFC, PCNA, dATP, and dCTP were introduced, before template unwinding and DNA synthesis were initiated by addition of ATP, dTTP, dGTP, and RPA. The reaction was performed in parallel on a non-chromatinized DNA substrate. Denaturing gel electrophoresis of the fluorescently labeled DNA products of leading strand synthesis by Pol ε revealed two products unique to the nucleosomal substrate (Extended Data Fig. 1g), indicating that the nucleosome is a barrier to replisome progression.

To gain structural insight into replisome traversal of the nucleosome, we performed a replication reaction on a nucleosomal substrate (Methods, Extended Data Fig. 1b,c) and purified replisome-nucleosome complexes by gradient ultracentrifugation with mild cross-linking (Extended Data Fig. 1h-k). Single-particle cryo-EM analysis of the sample revealed a replisome at an overall resolution of 3.3 Å with diffuse density downstream of the parental DNA (Extended Data Fig. 2, Table 1). Signal subtraction and classification of this density resulted in 2D classes resembling nucleosomal side views (Extended Data Fig. 2a). The low number of particles, however, prevented high-resolution 3D reconstruction of the nucleosome.

**Table 1.** Cryo-EM data collection statistics for dataset 1.

|  |  |
| --- | --- |
| Microscope | Thermo Fisher Titan Krios |
| Voltage (keV) | 300 |
| Camera | Falcon IV |
| Magnification | 105,000 |
| Pixel size at detector (Å per pixel) | 1.19 |
| Total electron exposure (e<sup>-</sup> Å<sup>-2</sup>) | 53.41 |
| Exposure rate (e<sup>-</sup> Å<sup>-2</sup> s<sup>-1</sup>) | 7.87 |
| Number of frames collected during exposure | 58 |
| Automation software | EPU |
| Micrographs collected (no.) | 7,014 |
| Micrographs used (no.) | 7,014 |
| Total extracted particles (no.) | 1,271,152 |
|  | Replisome |
|  | (Map 1) |
|  | EMD- |
| Final particles (no.) | 178,356 |
| Resolution (global, Å) |  |
| FSC 0.143 masked | 3.3 |
| FSC 0.5 masked | 3.7 |
| Map sharpening *B* factor (Å<sup>2</sup>) | 70.1 |
| 3DFSC sphericity | 0.954 |
| Map sharpening methods | cryoSPARC |

### Cryo-EM structures of replisome-nucleosome complexes

To facilitate the structural characterization of replisome-nucleosome engagement, we prepared a modified substrate, in which the double-stranded DNA in between the fork junction and nucleosome entry site was shortened to 22 bp (Extended Data Fig. 3a). The length was chosen to be short enough to capture the initial interaction between the replisome and the nucleosome before CMG translocation, but long enough to prevent nucleosome disruption by the pre-loaded replisome. A replisome consisting of CMG, TIMELESS-TIPIN, AND-1, and CLASPIN was assembled on the forked substrate and translocation into the nucleosome was initiated by addition of ATP (Methods). The resulting replisome-nucleosome complexes were purified by gradient ultracentrifugation with mild cross-linking (Extended Data Fig. 3) and analyzed by single-particle cryo-EM, yielding four structures of the replisome engaging the nucleosome in distinct conformational states (Fig. 1, Extended Data Figs. 4-6, Table 2, and Supplementary Video 1). After initial cryo-EM analysis, we obtained a replisome reconstruction at 2.1 Å. Further classifications and refinements revealed the four distinct replisome-nucleosome complexes. The nucleosomes in these structures were resolved at 2.8 Å, 3.0 Å, 7.8 Å, and 7.8 Å, with clear side-chain densities or secondary structure elements (Extended Data Figs. 5-7).

**Figure 1.**
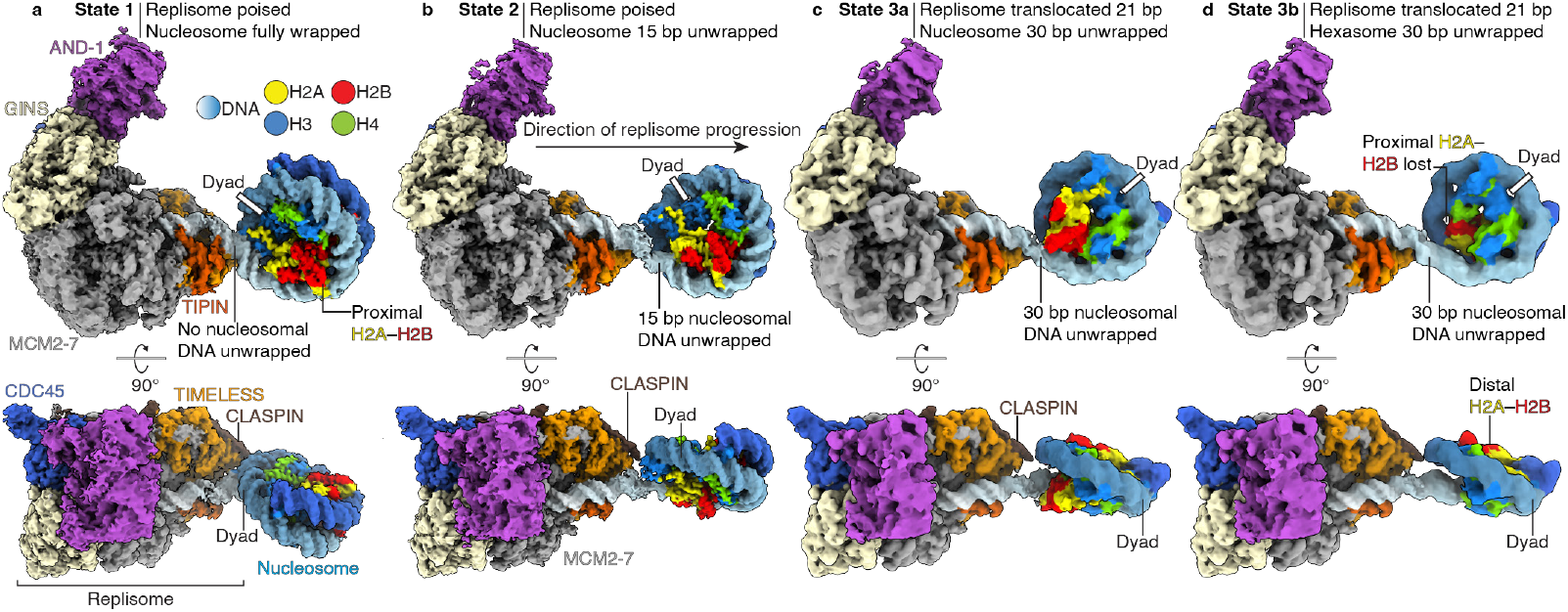
Cryo-EM structures of replisome progression into a nucleosome. **a**, Structure of the replisome engaging a fully wrapped nucleosome (state 1). **b**, Structure of the replisome engaging a 15 bp unwrapped nucleosome (state 2). **c**, Structure of the replisome engaging a 30 bp unwrapped nucleosome (state 3a). **d**, Structure of the replisome engaging a 30 bp unwrapped hexasome with the proximal H2A–H2B dimer lost (state 3b). Color code used throughout. Coulomb potential maps shown for all structures.

**Table 2.**
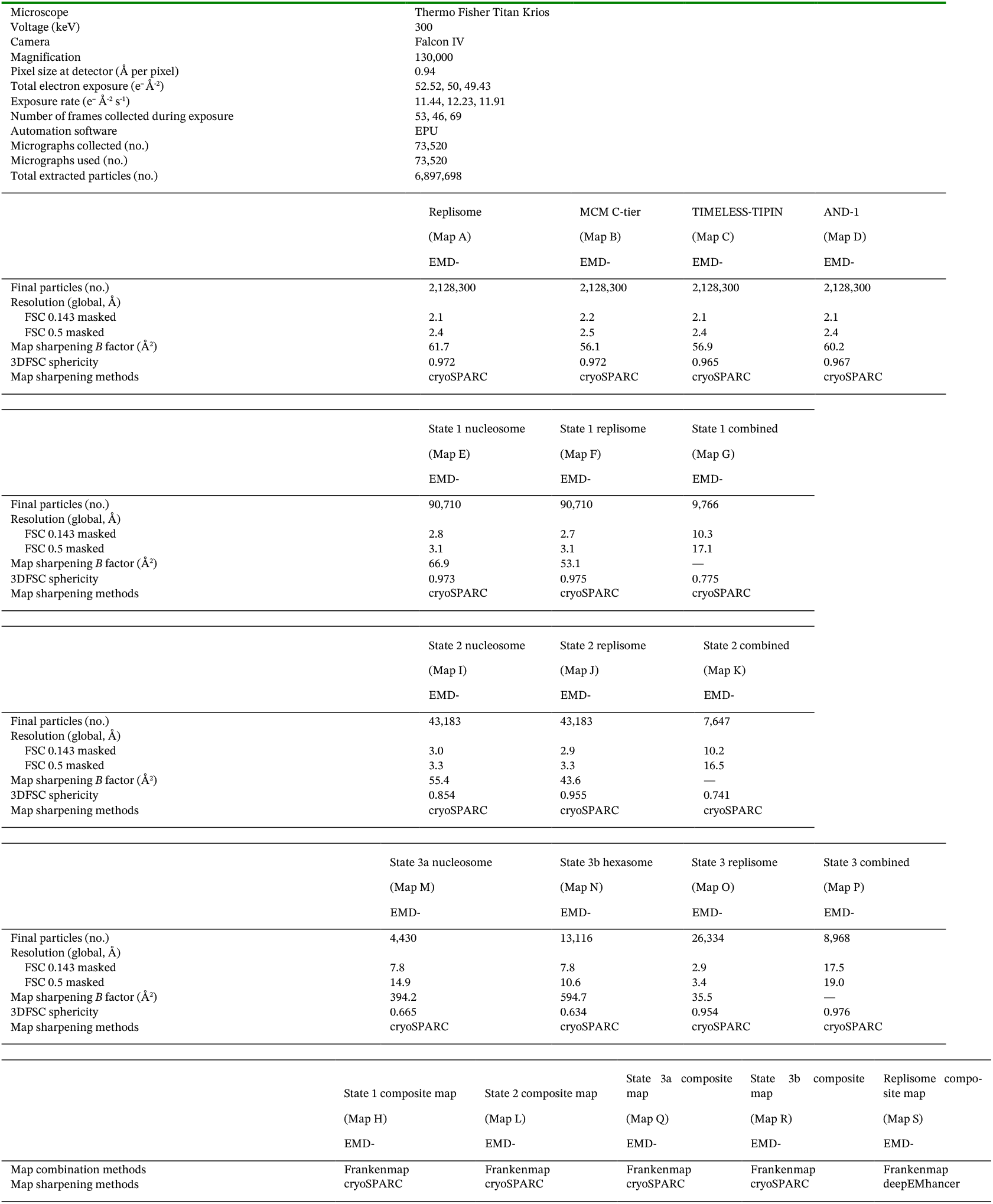
Cryo-EM data collection statistics for datasets A, B, and C.

|  |  |
| --- | --- |
| Microscope | Thermo Fisher Titan Krios |
| Voltage (keV) | 300 |
| Camera | Falcon IV |
| Magnification | 130,000 |
| Pixel size at detector (Å per pixel) | 0.94 |
| Total electron exposure (e<sup>-</sup> Å<sup>-2</sup>) | 52.52, 50, 49.43 |
| Exposure rate (e<sup>-</sup> Å<sup>-2</sup> s<sup>-1</sup>) | 11.44, 12.23, 11.91 |
| Number of frames collected during exposure | 53, 46, 69 |
| Automation software | EPU |
| Micrographs collected (no.) | 73,520 |
| Micrographs used (no.) | 73,520 |
| Total extracted particles (no.) | 6,897,698 |

|  |  |  |  |  |
| --- | --- | --- | --- | --- |
|  | Replisome | MCM C-tier | TIMELESS-TIPIN | AND-1 |
|  | (Map A) | (Map B) | (Map C) | (Map D) |
|  | EMD- | EMD- | EMD- | EMD- |
| Final particles (no.) | 2,128,300 | 2,128,300 | 2,128,300 | 2,128,300 |
| Resolution (global, Å) |  |  |  |  |
| FSC 0.143 masked | 2.1 | 2.2 | 2.1 | 2.1 |
| FSC 0.5 masked | 2.4 | 2.5 | 2.4 | 2.4 |
| Map sharpening *B* factor (Å<sup>2</sup>) | 61.7 | 56.1 | 56.9 | 60.2 |
| 3DFSC sphericity | 0.972 | 0.972 | 0.965 | 0.967 |
| Map sharpening methods | cryoSPARC | cryoSPARC | cryoSPARC | cryoSPARC |

|  |  |  |  |
| --- | --- | --- | --- |
|  | State 1 nucleosome | State 1 replisome | State 1 combined |
|  | (Map E) | (Map F) | (Map G) |
|  | EMD- | EMD- | EMD- |
| Final particles (no.) | 90,710 | 90,710 | 9,766 |
| Resolution (global, Å) |  |  |  |
| FSC 0.143 masked | 2.8 | 2.7 | 10.3 |
| FSC 0.5 masked | 3.1 | 3.1 | 17.1 |
| Map sharpening *B* factor (Å<sup>2</sup>) | 66.9 | 53.1 | — |
| 3DFSC sphericity | 0.973 | 0.975 | 0.775 |
| Map sharpening methods | cryoSPARC | cryoSPARC | cryoSPARC |
|  | State 2 nucleosome | State 2 replisome | State 2 combined |
|  | (Map I) | (Map J) | (Map K) |
|  | EMD- | EMD- | EMD- |
| Final particles (no.) | 43,183 | 43,183 | 7,647 |
| Resolution (global, Å) |  |  |  |
| FSC 0.143 masked | 3.0 | 2.9 | 10.2 |
| FSC 0.5 masked | 3.3 | 3.3 | 16.5 |
| Map sharpening *B* factor (Å<sup>2</sup>) | 55.4 | 43.6 | — |
| 3DFSC sphericity | 0.854 | 0.955 | 0.741 |
| Map sharpening methods | cryoSPARC | cryoSPARC | cryoSPARC |

|  |  |  |  |  |
| --- | --- | --- | --- | --- |
|  | State 3a nucleosome | State 3b hexasome | State 3 replisome | State 3 combined |
|  | (Map M) | (Map N) | (Map O) | (Map P) |
|  | EMD- | EMD- | EMD- | EMD- |
| Final particles (no.) | 4,430 | 13,116 | 26,334 | 8,968 |
| Resolution (global, Å) |  |  |  |  |
| FSC 0.143 masked | 7.8 | 7.8 | 2.9 | 17.5 |
| FSC 0.5 masked | 14.9 | 10.6 | 3.4 | 19.0 |
| Map sharpening *B* factor (Å<sup>2</sup>) | 394.2 | 594.7 | 35.5 | — |
| 3DFSC sphericity | 0.665 | 0.634 | 0.954 | 0.976 |
| Map sharpening methods | cryoSPARC | cryoSPARC | cryoSPARC | cryoSPARC |

|  |  |  |  |  |  |
| --- | --- | --- | --- | --- | --- |
|  | State 1 composite map | State 2 composite map | State 3a composite map | State 3b composite map | Replisome composite map |
|  | (Map H) | (Map L) | (Map Q) | (Map R) | (Map S) |
|  | EMD- | EMD- | EMD- | EMD- | EMD- |
| Map combination methods | Frankenmap | Frankenmap | Frankenmap | Frankenmap | Frankenmap |
| Map sharpening methods | cryoSPARC | cryoSPARC | cryoSPARC | cryoSPARC | deepEMhancer |

Generation of composite maps of the replisome-nucleosome complexes allowed unambiguous docking of replisome^10^ and nucleosome^11^ structures into the cryo-EM reconstructions (Methods). The refined pseudo-atomic models showed good stereochemistry (Tables 3 and 4).

**Table 3.** Model composition, refinement, and validation.

|  | Replisome | Replisome-nucleosome | Replisome-nucleosome | Replisome-nucleosome | Replisome-nucleosome |
| --- | --- | --- | --- | --- | --- |
|  |  | State 1 | State 2 | State 3a | State 3b |
|  | PDB: | PDB: | PDB: | PDB: | PDB: |
|  | EMD- | EMD- | EMD- | EMD- | EMD- |
| **Model composition** |  |  |  |  |  |
| Non-hydrogen atoms (no.) | 58,632 | 70,531 | 70,534 | 69,674 | 68,153 |
| Protein residues (no.) | 7,176 | 7,927 | 7,927 | 7,925 | 7,729 |
| Ligands (no.) | 12 | 12 | 12 | 12 | 12 |
| Nucleotide residues (no.) | 57 | 347 | 347 | 305 | 305 |
| **Model Refinement** |  |  |  |  |  |
| Initial models used (PDB #) | 7PLO | 7PLO, 3LZ0 | 7PLO, 3LZ0 | 7PLO, 3LZ0 | 7PLO, 3LZ0 |
| Refinement packages | Coot, PHENIX | Coot, PHENIX | Coot, PHENIX | Coot, PHENIX | Coot, PHENIX |
| Model-Map scores |  |  |  |  |  |
| Cross-correlation coefficient | 0.84 | 0.71 | 0.69 | 0.54 | 0.55 |
| Model resolution (Å) | 1.5 | 2.8 | 3.0 | 3.2 | 3.1 |
| FSC threshold | 0.143 | 0.143 | 0.143 | 0.143 | 0.143 |
| Mean *B* factors (Å²) |  |  |  |  |  |
| Protein residues (no.) | 123.29 | 123.29 | 123.29 | 123.29 | 123.29 |
| Ligands (no.) | 137.86 | 137.86 | 137.86 | 137.86 | 137.86 |
| Nucleotide residues (no.) | 145.47 | 176.03 | 198.78 | 176.03 | 176.03 |
| R.m.s. deviations from ideal values |  |  |  |  |  |
| Bond lengths (Å) | 0.005 | 0.006 | 0.007 | 0.009 | 0.009 |
| Bond angles (°) | 1.174 | 1.267 | 1.313 | 1.295 | 1.290 |
| **Validation** |  |  |  |  |  |
| MolProbity score | 1.71 | 1.97 | 2.03 | 2.00 | 1.92 |
| CaBLAM outliers (%) | 1.94 | 1.87 | 1.88 | 1.89 | 1.89 |
| Clashscore | 5.61 | 6.68 | 7.67 | 6.99 | 6.64 |
| Poor rotamers (%) | 1.62 | 2.43 | 2.45 | 2.46 | 2.19 |
| C-beta deviations (%) | 0.00 | 0.03 | 0.03 | 0.03 | 0.03 |
| EMRinger score | 3.68 | 3.15 | 2.78 | 1.57 | 1.64 |
| Ramachandran plot |  |  |  |  |  |
| Favored (%) | 96.34 | 95.50 | 95.50 | 95.51 | 95.73 |
| Allowed (%) | 3.41 | 4.11 | 4.11 | 4.10 | 3.87 |
| Outliers (%) | 0.25 | 0.40 | 0.40 | 0.40 | 0.39 |

**Table 4.**
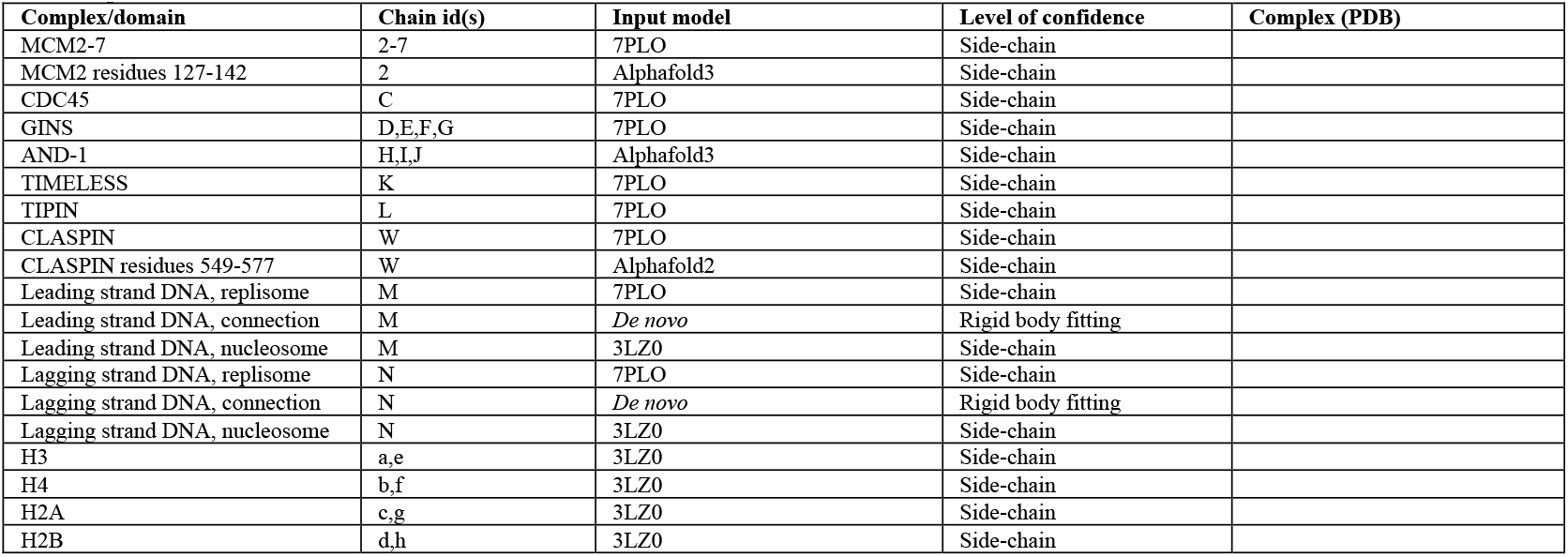
Input structural models and model confidence.

Compared to previous structures, we resolved additional replisomal elements. First, we identified a two-stranded β-sheet in CLASPIN (residues 549-577, Extended Data Fig. 7w) that interacts with the C-tier of MCM2 and potentially stabilizes the MCM2 winged-helix domain. Second, we revealed a divalent cation that coordinates histidine 615 from each AND-1 copy, facilitating AND-1 trimerization (Extended Data Fig. 7x). Third, our reconstructions facilitated assignment of four nucleotides in the MCM5:3, MCM2:5, MCM6:2 and MCM4:6 ATPase sites (Methods, Extended Data Fig. 7y). Additionally, a number of previously unmodelled flexible loops across the replisome could be sufficiently resolved to build confident pseudo-atomic models for these elements. The replisome maintained similar conformations in all four states (Fig. 1).

### Nucleosome transitions during replisome progression

Our structures captured nucleosome transitions during replisome progression with four distinct conformational states: one fully wrapped (state 1) and three partially unwrapped (states 2, 3a and 3b) (Figs. 1 and 2). In state 1, the nucleosome remained fully wrapped, positioned directly downstream of the poised, non-translocated replisome (Fig. 2a). In state 2, the nucleosome was unwrapped by 15 bp (from superhelical location (SHL) –7 to –5.5), positioning the unwrapped DNA for entry into the poised replisome (Fig. 2b). In states 3a and 3b, the replisome had translocated 21 bp from its original position and entered the nucleosome, with nucleosomal DNA unwrapped from SHL –7 to –4 (Fig. 2c,d). In state 3a, the histone octamer remained intact; in contrast, state 3b showed loss of the proximal H2A–H2B dimer, resulting in a hexasome. The loss of the dimer is likely due to its destabilization following DNA unwrapping. The degree of nucleosome unwrapping in our structures corresponds to sites of histone-DNA contacts^4^.

**Figure 2.**
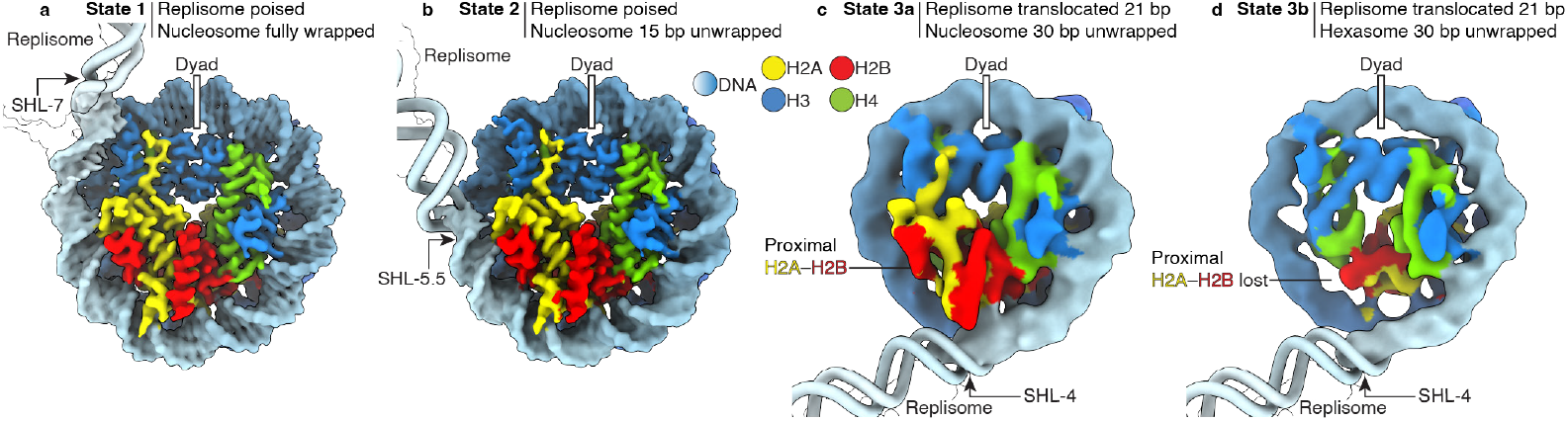
Comparison of fully wrapped and partially unwrapped nucleosomes and hexasome. **a**, Structure of the fully wrapped nucleosome (state 1). **b**, Structure of the 15 bp unwrapped nucleosome (state 2). **c**, Structure of the 30 bp unwrapped nucleosome (state 3a). **d**, Structure of the 30 bp unwrapped hexasome with the proximal H2A–H2B dimer lost (state 3b). Coulomb potential maps shown for nucleosomes and hexasome. Fitted models shown for replisome and additional DNA.

In all states, the nucleosomes remained positioned immediately downstream of the leading edge of TIMELESS-TIPIN, indicating that this is the site of parental nucleosome unwrapping during replication. Accordingly, the downstream DNA exhibited a pronounced downward kink at the TIMELESS-TIPIN-nucleosome interface (Extended Data Fig. 8a-c). In contrast, a later intermediate of replisomal histone transfer in yeast, in which the nucleosome was fully disassembled, displayed a linear DNA trajectory downstream of the replisome^12^ (Extended Data Fig. 8d). The orientation of the nucleosomal disk relative to the replisome remained largely similar across the four states.

The loss of the proximal H2A–H2B dimer in state 3b demonstrates that the unwrapping of nucleosomal DNA by the progressing replisome can disrupt nucleosomal integrity. A similar displacement of the proximal H2A– H2B dimer has been observed in transcription through nucleosomes^13,14^ (Extended Data Fig. 8e,f). Importantly, nucleosome disruption by the approaching replisome rationalizes the requirement for histone chaperones during DNA replication, particularly the histone chaperone FAcilitates Chromatin Transcription (FACT)^5,12,15,16^. Superposition of FACT onto the nucleosome (state 3a) shows that sufficient proximal H2A–H2B dimer surface is exposed to allow for FACT binding via the SPT16 C-terminal domain^17^ (Extended Data Fig. 8g,h). This mechanism mirrors nucleosome transcription intermediates in which FACT assists transcription elongation complexes^14,18^, preventing histone dissociation. Taken together, the structural basis of replisome-nucleosome engagement has striking parallels with transcription-nucleosome engagement.

### Model for replisome passage through a nucleosome

Our structures, combined with previous biochemical and structural data^12,16,19–22^, suggest a stepwise mechanism for nucleosome engagement and DNA unwrapping during replication (Extended Data Fig. 8e). First, the encroaching replisome unwraps DNA from the parental nucleosome. Further unwrapping of nucleosomal DNA exposes the replisome-proximal H2A–H2B dimer, enabling the engagement of FACT. Next, progression of the replisome beyond the dyad results in removal of the histones from the parental DNA and initial storage of a FACT-histone hexamer intermediate by MCM2 and TIME-LESS on the replisome^12^. Replisome factors including CLASPIN^16,19,20^, Pol ε^21^, and Pol α^22^ co-chaperone the histones across the replisome surface. Finally, the histones with their associated epigenetic information are deposited on one of the two newly synthesized DNA strands. Going forward, our *in vitro* reconstituted chromatin replication system and replisome-nucleosome structures lay the foundation to elucidate the structural basis of epigenetic inheritance by investigating the mechanisms of replication-coupled histone transfer and nucleosome (re)assembly.

## Methods

No statistical methods were used to predetermine sample size. The experiments were not randomized, and the investigators were not blinded to allocation during experiments and outcome assessment.

### Cloning and protein expression

*H. sapiens* MCM3, MCM5, MCM6, PSF1, PSF2, PSF3, SLD5 and CDC45 were amplified from cDNA and cloned into vector 438-A by ligation-independent cloning (LIC). *H. sapiens* MCM2 and MCM7 were amplified from Addgene plasmid #116949 (a gift from James Chong) and cloned into vector 438-A by LIC. *H. sapiens* MCM4 was amplified from DNASU clone HsCD00859736 and cloned into vector 438-A by LIC. CDC45 was encoded with an internal FLAG tag and SLD5 with an N-terminal Twin-Strep tag as described^7^. The internal FLAG tag was introduced into 438-A-CDC45 by ‘Round-The-Horn mutagenesis and the N-terminal Twin-Strep tag was introduced into 438-A-SLD5 by sequence- and ligation-independent cloning (SLIC). MCM2-7 and GINS (PSF1, PSF2, PSF3 and SLD5) were combined into vectors pBIG2ab and pBIG1a, respectively^23^. Bacmid, virus, and protein production for CMG were performed as described^24^.

*H. sapiens* TIMELESS and TIPIN were amplified from cDNA and cloned into vectors 438-C and 438-A, respectively. Using LIC, the two genes were combined on a single 438-series vector. The construct contained TIME-LESS with an N-terminal His6 tag, followed by a maltose binding protein (MBP) tag, and a TEV protease cleavage site. TIPIN did not contain a tag. *H. sapiens* AND-1 was amplified from DNASU clone HsCD00868051 and cloned into vector 438-C, containing an N-terminal TEV protease cleavage site, followed by a maltose binding protein (MBP) tag, and a His6 tag, by LIC. *H. sapiens* CLASPIN was amplified from cDNA and cloned into a 438-series vector containing a C-terminal TEV protease cleavage site, followed by a maltose binding protein (MBP) tag, and a His6 tag, by LIC. Bacmid, virus, and protein production for TIMELESS-TIPIN, AND-1 and CLASPIN were performed as described^24^, except that CLASPIN was expressed in Sf21 insect cells.

*H. sapiens* POLE1 was amplified from DNASU clone HsCD00860166 and cloned into vector 438-A by LIC. *H. sapiens* POLE2 was amplified from cDNA and cloned into vector 438-A by LIC. *H. sapiens* POLE3 was ordered as a gBlock (Integrated DNA Technologies) and cloned into vector 438-C, containing an N-terminal TEV protease cleavage site, followed by a maltose binding protein (MBP) tag, and a His6 tag, by LIC. *H. sapiens* POLE4 was ordered as a gBlock (Integrated DNA Technologies) and cloned into vector 438-C by LIC. POLE1-4 were combined into vector pBIG2ab^23^. Bacmid, virus, and protein production for Pol ε were performed as described^24^.

*H. sapiens* RFC1 was amplified from DNASU clone HsCD00852551 and cloned into vector 438-C, containing an N-terminal TEV protease cleavage site, followed by a maltose binding protein (MBP) tag, and a His6 tag, by LIC. *H. sapiens* RFC2 was was amplified from DNASU clone HsCD00829507 and cloned into vector 438-A by LIC. *H. sapiens* RFC3, RFC4 and RFC5 were amplified from cDNA and cloned into vector 438-A by LIC. RFC1-5 were combined into vector pBIG2ab^23^. Bacmid, virus, and protein production for RFC were performed as described^24^.

*H. sapiens* PCNA was expressed as described^7^ from Addgene plasmid #134898 (a gift from Andrew Deans). *H. sapiens* RPA was expressed as described^7^ from Addgene plasmid #102613 (a gift from Marc Wold).

### Protein purification

*H. sapiens* CMG was purified as described^7^, except that a UNO Q Polishing Column #7200009 (Bio-Rad) was used in place of the MonoQ PC 1.6/5 (GE Healthcare). *H. sapiens* PCNA and RPA were purified as described^7^. *X. laevis* histones were purified as described^24^.

*H. sapiens* TIMELESS-TIPIN, AND-1, CLASPIN, Pol ε and RFC were purified as follows. TIMELESS-TIPIN, AND-1, Pol ε and RFC were each purified from 600 mL Hi5 cells. CLASPIN was purified from 1.2 L Sf21 cells. Protein purifications were performed at 4 °C. Cells were resuspended in lysis buffer (300 mM NaCl, 20 mM Na·HEPES pH 7.4 at 25 °C, 10% (v/v) glycerol, 30 mM imidazole pH 8, 1 mM TCEP pH 8, 0.284 µg ml^−1^ leupeptin, 1.37 µg ml^−1^ pepstatin A, 0.17 mg ml^−1^ PMSF and 0.33 mg ml^−1^ benzamidine) and lysed by sonication. Lysed cells were subsequently centrifuged and then ultra-centri-fuged. The supernatant was cleared through a 0.45 µM filter and loaded onto a 5 mL HisTrap HP (Cytiva) column equilibrated in lysis buffer. After 10 CV washes with lysis buffer, a self-packed XK column (Cytiva) with 15 mL of Amylose resin (New England Biolabs) was attached to the HisTrap column. Sample was directly eluted onto the Amylose column with 5 CV nickel elution buffer (300 mM NaCl, 20 mM Na·HEPES pH 7.4 at 25 °C, 10% (v/v) glyc-erol, 500 mM imidazole pH 8, 1 mM TCEP pH 8). The HisTrap column was then removed, and the Amylose column was washed with 5 CV lysis buffer. Sample was eluted with 5 CV amylose elution buffer (300 mM NaCl, 20 mM Na·HEPES pH 7.4 at 25 °C, 10% (v/v) glycerol, 30 mM imidazole pH 8, 116.9 mM maltose, 1 mM TCEP pH 8). Peak fractions were pooled, mixed with 1.5 mg of TEV protease with an N-terminal MBP tag, and dialysed overnight in 7 kDa MWCO SnakeSkin dialysis tubing (Thermo Scientific) against dialysis buffer (150 mM NaCl, 20 mM Na·HEPES pH 7.4 at 25 °C, 10% (v/v) glycerol, 1 mM TCEP pH 8). The sample was then applied to a self-packed XK column with 15 mL of Amylose resin in tandem with a 5 mL HiTrap Q (Cytiva) column, both equilibrated in dialysis buffer.

The columns were washed with 5 CV of dialysis buffer after which the Amylose column was removed. The protein was eluted from the HiTrap Q column with a linear gradient from dialysis buffer to high salt buffer (1 M NaCl, 20 mM Na·HEPES pH 7.4 at 25 °C, 10% (v/v) glycerol, 1 mM TCEP pH 8) over 25 CV. Peak fractions were concentrated using an Amicon 30,000 MWCO centrifugal filter unit (Millipore). The concentrated sample was applied to a Superose 6 Increase 10/300 GL (Cytiva) equilibrated in gel filtration buffer (300 mM NaCl, 20 mM Na·HEPES pH 7.4 at 25 °C, 10% (v/v) glycerol, 1 mM TCEP pH 8). The elution was fractionated and analyzed by SDS-PAGE. Peak fractions were concentrated with an Amicon 30,000 MWCO centrifugal filter unit, aliquots were frozen in liquid nitrogen, and stored at −80 °C.

### Helicase assays

Helicase assays were performed as described^7^ except that CMG was loaded onto the substrate for 90 min and reactions were run on self-cast 10 % TBE polyacryla-mide gels.

### Preparation of substrates A and B

A vector containing a modified Widom 601 sequence was used as a template for PCR. Large-scale PCR reactions were performed with PCR primers (forward primer: 5′-ACG AAG CGT AGC ATC ACT GTC TTG-3′; reverse primers: 5′-ATC AGA ATC CCG GTG CCG AG-3′ for substrate A and 5′-AGT ATT GAG CCT CAG GAA ACA GCT ATG-3′ for substrate B) at a scale of 50 ml, and resulted in PCR products 5′-ACG AAG CGT AGC ATC ACT GTC TTG TGT TTG GTG TGT CAA GGT GGT GGC CGT TTT CGA TGT ATA TAT CTG ACA CGT GCC TGG AGA CTA GGG AGT AAT CCC CTT GGC GGT TAA AAC GCG GGG GAC AGC GCG TAC GTG CGT TTA AGC GGT GCT AGA GCT GTC TAC GAC CAA TTG AGC GGC CTC GGC ACC GGG ATT CTG AT-3′ (top strand) for substrate A and 5′-ACG AAG CGT AGC ATC ACT GTC TTG TGT TTG GTG TGT CAA GGT GGT GGC CGT TTT CGA TGT ATA TAT CTG ACA CGT GCC TGG AGA CTA GGG AGT AAT CCC CTT GGC GGT TAA AAC GCG GGG GAC AGC GCG TAC GTG CGT TTA AGC GGT GCT AGA GCT GTC TAC GAC CAA TTG AGC GGC CTC GGC ACC GGG ATT CTG ATA TCG CGC GTG ATC TTA CGG CAT TAT ACG TAT GAT CGG TCC ACG ATC AGC TAG ATT ATC TAG TCA GCT TGA TGT CAT AGC TGT TTC CTG AGG CTC AAT ACT-3′ (top strand) for substrate B. Each PCR product was purified using anion exchange chromatography (Cytiva 5 mL HiTrap Q) followed by ethanol precipitation. The DNA product was digested with TspRI in 1X CutSmart Buffer (NEB) overnight at 65 °C to generate the 9 nt single-stranded DNA overhang. The digestion product was purified using a Model 491 Prep Cell (Bio-Rad), ethanol pre-cipitated and resuspended in water. DNA Oligos 5′-ACC AAA CAC AAG ACA GTG ATG CTA TGT GGT AGG AAG TGA GAA TTG GAG AGT TTT TTT TTT TTT TTT TTT TTT TTT TTT TTT TTT TTT TTT TGA GGA AAG AAT GTT GGT GAG GGT TGG GAA GTG GAA GGA TGG GCT CGA GAG GTT TTT TTT TTT TTT TTT TTT TTT TTT TTT TTT TT-3′ and 5′-TTT TTT TTT TTT TTT TTT TTA CTC TCC AAT TCT CAC TTC CTA CCA CAT AGC ATC ACT GTC TTG TGT TTG GT-3′ (Integrated DNA Technologies) were mixed at 50 µM in a buffer containing 50 mM NaCl, 20 mM Tris-HCl pH 7.5 and 0.5 mM EDTA, and annealed by incubating at 95 °C for 5 min and then decreasing the temperature by 1 °C min^−1^ steps to a final temperature of 4 °C. The resulting forked DNA was digested with TspRI in 1X CutSmart Buffer (NEB) at 65 °C overnight to generate the complementary 9 nt single-stranded DNA overhang, purified using a Model 491 Prep Cell (Bio-Rad), ethanol precipitated and resuspended in water. A 2-fold molar excess of the TspRI-digested fork was mixed with 0.5 µM TspRI-digested PCR product in 100 µL reactions each containing 2000 units of T4 DNA ligase (NEB) and 1X T4 DNA ligase buffer and incubated at 16 °C overnight. The DNA was extracted with phenol:chloroform:isoamyl alcohol 25:24:1 (Sigma-Aldrich), ethanol precipitated and resuspended in water, and purified using a Model 491 Prep Cell to remove excess un-ligated fork. Nucleosome core particle reconstitution was then performed using the salt-gradient dialysis method^25^.

### DNA replication assays

Replication reactions were set up at 37 °C in a buffer containing 100 mM potassium glutamate, 25 mM Na·HEPES pH 7.4, 0.005 % Tween-20, 1 mM DTT, 2 mM Mg(OAc)^2^ and 0.1 mg ml^−1^ BSA^9^. All concentrations refer to final reaction conditions. 100 nM CMG was pre-incubated with 50 nM DNA or nucleosomal substrate for 90 min in reconstitution buffer supplemented with 0.1 mM AMP-PNP. 50 nM DNA primer (5’-/6-FAM/-CCT CTC GAG CCC ATC CTT CCA CTT CCC AAC CCT CAC C-3’) was added and the reaction was incubated for 5 min. TIMELESS–TIPIN, AND-1, CLASPIN, Pol ε, RFC, PCNA (100 nM), dATP and dCTP (30 µM) were added and the reaction was incubated for 4 min. RPA (200 nM), dTTP, dGTP (30 µM) and ATP (4 mM) were added to initiate template unwinding and DNA synthesis. At specified time-points, 2 μL reactions were quenched with 1 μL STOP buffer (200 mM Tris-HCl pH 8.0, 0.5 mg/mL Proteinase K (Roche), and 80 mM EDTA) and incubated at 25 °C for 10 min. 12 μL formamide (Sigma-Aldrich) was added to each sample, and the samples were denatured at 95 °C for 10 minutes before being separated by a 9 % denaturing urea gel (8 M urea, 1X TBE, 12 % acrylamide:bis-acrylamide, 10 % formamide). DNA products were detected using a 6-FAM fluorescence scan on a Typhoon 5 with an automatically determined PMT. DNA replication assays were performed as two independent replicates.

### CMG-TIMELESS-TIPIN-AND-1-CLASPIN-Pol ε-RPA-substrate B complex preparation for cryo-EM

Glycerol gradient preparation was performed as described^7^. The reconstitution reaction was set up at 37 °C in a buffer containing 100 mM potassium glutamate, 25 mM Na·HEPES pH 7.4, 0.005 % Tween-20, 1 mM DTT, 2 mM Mg(OAc)^2^ and 0.1 mg ml^−1^ BSA, to yield a final volume of 550 µL. All concentrations refer to final reaction conditions. 100 nM CMG was pre-incubated with 150 nM nucleosomal substrate for 90 min in reconstitution buffer supplemented with 0.1 mM AMP-PNP. 150 nM DNA primer (5’-/6-FAM/-CCT CTC GAG CCC ATC CTT CCA CTT CCC AAC CCT CAC C-3’) was added and the reaction was incubated for 5 min. TIMELESS– TIPIN, AND-1, CLASPIN, Pol ε (150 nM), dATP and dCTP (30 µM) were added and the reaction was incubated for 4 min. RPA (200 nM), dTTP, dGTP (30 µM) and ATP (4 mM) were added to initiate template unwinding and DNA synthesis. The reaction was incubated for 15 min at which point 0.5 mM AMP-PNP was added and the sample was transferred to 4 °C. ∼180 μL of the sample was loaded onto each gradient: one lacking and two containing crosslinker. Gradient centrifugation and fractionation were performed as described^**7**^. Silver-stained SDS–PAGE of fractions with and without cross-linkers, and denaturing urea PAGE of fractions without crosslinkers, were used to identify fractions containing replisome-nucleosome complexes. Buffer exchange of selected, pooled fractions was performed as described^**7**^. Quantifoil R 3.5/1, 200 Mesh, Copper cryo-EM grids (Electron Microscopy Sciences) pre-coated with an ultrathin (3–5 nm) amorphous carbon were glow discharged for 30 s at 15 mA using a Pelco Easiglow plasma discharge system. 3 μL of sample was applied to the top, carbon-coated side of the grid. After incubation for 45 s and blotting for 8 s, the grid was vitrified by plunging it into liquid ethane using a Vitrobot Mark IV (FEI) operated at 4 °C and 100 % humidity.

### Preparation of substrate C

Ultramer DNA Oligos (Integrated DNA Technologies) 5′-ATC AGA ATC CCG GTG CCG AGG CCG CTC AAT TGG TCG TAG ACA GCT CTA GCA CCG CTT AAA CGC ACG TAC GCG CTG TCC CCC GCG TTT TAA CCG CCA AGG GGA TTA CTC CCT AGT CTC CAG GCA CGT GTC AGA TAT ATA CAT CGA AAT TAG AGA ATT GGA GAG TGT GTT TTT TTT TTT TTT TTT TTT TTT TTT TTT TTT TT-3′ and 5′-GGC AGG CAG GCA GGC ACA CAC TCT CCA ATT CTC TAA TTT CGA TGT ATA TAT CTG ACA CGT GCC TGG AGA CTA GGG AGT AAT CCC CTT GGC GGT TAA AAC GCG GGG GAC AGC GCG TAC GTG CGT TTA AGC GGT GCT AGA GCT GTC TAC GAC CAA TTG AGC GGC CTC GGC ACC GGG ATT CTG AT-3′ were mixed at 50 µM in a buffer containing 50 mM NaCl, 20 mM Tris-HCl pH 7.5 and 0.5 mM EDTA, and annealed by incubating at 95 °C for 5 min and then decreasing the temperature by 1 °C min^−1^ steps to a final temperature of 4 °C in a thermocycler. The resulting forked DNA substrate was purified using a Model 491 Prep Cell and subsequently concentrated to ∼10 μM using an Amicon 10,000 MWCO centrifugal filter unit (Millipore). Subsequent nucleosome core particle reconstitution was performed using the salt-gradient dialysis method. The resulting nucleosome was purified using a Model 491 Prep Cell and concentrated to ∼2.5 μM using an Amicon 10,000 MWCO centrifugal filter unit. Quantification of the purified nucleosome was achieved by measuring absorbance at 280 nm. Molar extinction coefficients at 280 nm were determined for protein and nucleic acid components and were summed to yield a molar extinction coefficient for the reconstituted nucleosome substrate.

### CMG-TIMELESS-TIPIN-AND-1-CLASPIN-substrate C complex preparation for cryo-EM

Glycerol gradient preparation was performed as described^7^. The reconstitution reaction was set up at 37 °C in a final reconstitution buffer containing 100 mM potassium glutamate, 25 mM Na·HEPES pH 7.4, 0.005 % Tween-20, 1 mM DTT and 2 mM Mg(OAc)^2^, to yield a final volume of 550 µL. The concentrations of individual components in the final reaction were: 150 nM nucleosome, 100 nM CMG, 150 nM TIMELESS–TIPIN, 150 nM AND-1 and 150 nM CLASPIN. CMG (200 nM) was pre-incubated with the nucleosome (300 nM) for 100 min in reconstitution buffer supplemented with 0.1 mM AMP-PNP. The reaction was diluted 1.9-fold by the addition of reconstitution buffer containing TIMELESS–TIPIN, AND-1 and CLASPIN, and incubated for 10 min. 5 mM ATP was added to initiate DNA unwinding, bringing the reaction to its final volume of 550 µL. The reaction was incubated for 15 min, at which point 5 mM AMP-PNP was added and the sample was transferred to 4 °C. ∼200 μL of the sample was loaded onto each gradient: one lacking and two containing cross-linker. Gradient centrifugation, fractionation and buffer exchange of selected, pooled fractions were performed as described^7^. Quantifoil R 3.5/1, 200 Mesh, Copper cryo-EM grids (Electron Microscopy Sciences) pre-coated with an ultra-thin layer (3–5 nm) of amorphous carbon were glow discharged for 30 s at 15 mA using a Pelco Easiglow plasma discharge system. 3 μL of sample was applied to the carbon-coated side of the grid. After incubation for 45 s and blotting for 8 s, the grid was vitrified by plunging it into liquid ethane using a Vitrobot Mark IV (FEI) operated at 4 °C and 100 % humidity.

### Cryo-EM data acquisition

For the CMG-TIMELESS-TIPIN-AND-1-CLASPIN-Pol ε-RPA-substrate B complex, cryo-EM data was collected on a ThermoFisher Titan Krios at 300 keV equipped with a Falcon 4i direct electron detector, Selectris energy filter and Volta phase plate. Data collection was automated using Thermo Scientific Smart EPU software. The dataset was collected at a pixel size of 1.19 Å with a defocus range of 0.8 to 2 μm. The dataset yielded 7,014 micrographs. The dataset was collected with 58 movie frames at an exposure time of 6.79 s with an electron flux of 7.87 e^−^ per A^2^ per s for a total exposure dose of 53.41 e^−^ per Å^2^.

For the CMG-TIMELESS-TIPIN-AND-1-CLASPIN-substrate C complex, cryo-EM data were collected on a ThermoFisher Titan Krios at 300 keV equipped with a Falcon 4i direct electron detector, Selectris energy filter and Volta phase plate. Data collection was automated using Thermo Scientific Smart EPU software. Three datasets were collected at a pixel size of 0.94 Å with a defocus range of 0.8 to 2 μm. The first dataset yielded 27,351 micrographs. The second dataset yielded 19,088 micrographs. The third dataset yielded 27,081 micrographs. The first dataset was collected with 53 movie frames at an exposure time of 4.59 s with an electron flux of 11.44 e^−^ per A^2^ per s for a total exposure dose of 52.52 e^−^ per Å^2^. The second dataset was collected with 46 movie frames at an exposure time of 4.04 s with an electron flux of 12.23 e^−^ per A^2^ per s for a total exposure dose of 49.43 e^−^ per Å^2^. The third dataset was collected with 69 fractions at an exposure time of 4.04 s with an electron flux of 11.91 e^−^ per A^2^ per s for a total exposure dose of 48.11 e^−^ per Å^2^.

### Cryo-EM data analysis

For the CMG-TIMELESS-TIPIN-AND-1-CLASPIN-Pol ε-RPA-substrate B complex, initial image processing was conducted in cryoSPARC^26^ (v4.6.0). Movies were aligned using cryoSPARC Live Patch Motion Correction followed by CTF estimation. The cryoSPARC Blob picker was used to pick 1,271,152 particles. Particles were extracted at a box size of 490 pixels and down sampled to 250 pixels. Free nucleosome and junk particles were removed by 3 rounds of heterogeneous refinement, and the remaining 178,356 particles that revealed clear replisome density were re-extracted at a box size of 490 pixels with no downsampling and subjected to non-uniform refinement, resulting in a 3.3 Å map (map 1). Particle subtraction was performed in RELION-4.0^27^ on map 1 particles using a mask encompassing extra density downstream of the replisome. The subtracted particles were reimported into cryoSPARC and several rounds of heterogenous refinement and 2D classification were performed to remove junk particles, resulting in 3,338 particles in nucleosome-resembling 2D classes. The low number of particles in these classes prevented high-resolution 3D reconstruction of the nucleosome.

For the CMG-TIMELESS-TIPIN-AND-1-CLASPIN-substrate C complex, initial image processing was conducted in cryoSPARC. Movies were aligned using cryoSPARC Live Patch Motion Correction followed by CTF estimation. The cryoSPARC Blob picker was used to pick 6,897,698 particles. Particles were extracted at a box size of 500 pixels and down sampled to 250 pixels. Free nucleosome and junk particles were removed by 3 rounds of heterogeneous refinement, and the remaining 2,128,300 particles that revealed clear replisome density were re-extracted at a box size of 500 pixels with no downsampling and subjected to non-uniform refinement. Local and global CTF refinement and reference-based motion correction were then performed, followed by another round of non-uniform refinement, resulting in a 2.1 Å map (map A). Local refinement was performed using a mask encompassing the MCM C-tier, resulting in a 2.2 Å map (map B). Local refinement was performed using a mask encompassing TIMELESS-TIPIN, resulting in a 2.1 Å map (map C). Local refinement was performed using a mask encompassing AND-1, resulting in a 2.1 Å map (map D).

Particle subtraction was performed in RELION-4.0 on map A particles using a mask encompassing extra density downstream of the replisome. The subtracted particles were re-imported into cryoSPARC and subjected to three rounds of heterogeneous refinement, which revealed distinct hexasome and nucleosome classes. The hexasome class, which contained 221,439 particles, had DNA unwrapped to SHL –4. The nucleosome class, which contained 318,312 particles, was fully wrapped. The hexasome class was subjected to three further rounds of heterogeneous refinement to separate away remaining fully wrapped nucleosome particles, after which 55,949 hexasome particles remained. These were subjected to further rounds of heterogeneous refinement to remove remaining junk particles. The remaining particles were, in parallel, subjected to heterogeneous refinement and reverted to the original particles containing the replisome signal. The heterogeneous refinement revealed a distinct class of nucleosome particles, also unwrapped to SHL – 4, among the “hexasome” particles. The SHL –4 nucleosome and hexasome classes were each subjected to two further rounds of heterogeneous refinement to remove particles belonging to the other class, leaving 4,430 nucleosome particles and 13,116 hexasome particles. The particles in each class were subjected to non-uniform refinement, resulting in a 7.8 Å state 3a nucleosome map (map M) and a 7.8 Å state 3b hexasome map (map N).

The reverted SHL –4 nucleosome and hexasome particles containing the replisome signal were subjected to non-uniform refinement, resulting in a 3.0 Å replisome map (map O) with extra density downstream of the parental DNA. There appeared to be too much variability to simultaneously visualize both the complete replisome and the nucleosome. Therefore, another round of particle subtraction was performed in RELION-4.0 using a mask encompassing the MCM N-tier, TIMELESS-TIPIN, and the downstream extra density. The subtracted particles were re-imported into cryoSPARC and 2D classification was performed. Classes in which both replisome and nucleosome signal was visible were selected and subjected to heterogeneous refinement to remove poorly aligning particles, after which 8,968 particles remained. These were subjected to non-uniform refinement, resulting in a 17.5 Å state 3 combined replisome-nucleosome map (map P).

The fully wrapped nucleosome class containing 318,312 particles was subjected to further rounds of heterogeneous refinement, which revealed a distinct class of 15 bp unwrapped nucleosome particles among the “fully wrapped” nucleosome particles. The fully wrapped and 15 bp unwrapped classes were each subjected to two further rounds of heterogeneous refinement to remove particles belonging to the other class, leaving 90,710 fully wrapped nucleosome particles and 43,183 15 bp unwrapped nucleosome particles. The particles in each class were subjected to non-uniform refinement, resulting in a 2.8 Å state 1 nucleosome map (map E) and a 3.0 Å state 2 nucleosome map (map I).

The map E particles were reverted to the original particles containing the replisome signal and subjected to non-uniform refinement, resulting in a 2.7 Å replisome map (map F) with extra density downstream of the parental DNA. There appeared to be too much variability to simultaneously visualize both the complete replisome and the nucleosome. Therefore, another round of particle subtraction was performed in RELION-4.0 using a mask encompassing the MCM N-tier, TIMELESS-TIPIN, and the downstream extra density. The subtracted particles were re-imported into cryoSPARC and 2D classification was performed. Classes in which both replisome and nucleosome signal was visible were selected and subjected to heterogeneous refinement to remove poorly aligning particles, after which 9,766 particles remained. These were subjected to non-uniform refinement, resulting in a 10.3 Å state 1 combined replisome-nucleosome map (map G).

The same reversion and re-subtraction approach was taken for the map I state 2 nucleosome particles and resulted in a 2.9 Å replisome map (map J) and a 10.2 Å state 2 combined replisome-nucleosome map (map K) containing 7,647 particles.

The Frankenmap package in Warp^28^ and DeepEMhancer^29^ or the Sharpening Tools job in cryoSPARC were then used to produce the following composite maps. Maps B, C and D were placed into map A using the fitmap command and resampled using the vop resample command in UCSF ChimeraX^30^. Then, maps A, B, C and D were combined using Frankenmap and DeepEMhancer to produce the replisome composite map (map S).

Maps E and F were placed into map G using the fitmap command and resampled using the vop resample command in UCSF ChimeraX. Then, maps E and F were combined using Frankenmap and the Sharpening Tools job in cryoSPARC to produce the state 1 composite map (map H). Maps I and J were placed into map K using the fitmap command and resampled using the vop resample command in UCSF ChimeraX. Then, maps I, J and K were combined using Frankenmap and the Sharpening Tools job in cryoSPARC to produce the state 2 composite map (map L).

Maps M and O were placed into map P using the fitmap command and resampled using the vop resample command in UCSF ChimeraX. Then, maps M and O were combined using Frankenmap and the Sharpening Tools job in cryoSPARC to produce the state 3a composite map (map Q). Maps N and O were placed into map P using the fitmap command and resampled using the vop resample command in UCSF ChimeraX. Then, maps N and O were combined using Frankenmap and the Sharpening Tools job in cryoSPARC to produce the state 3b composite map (map R).

### Model building and refinement

CMG, TIMELESS-TIPIN, and CLASPIN from PDB 7PLO^10^ were rigid body docked into map A. AND-1 trimer was predicted using the AlphaFold3^31^ server and then rigid body docked into map D. DNA was manually built in Coot^32^. The resulting core replisome was manually adjusted in Coot using maps A, B, C, and D, and then real space refined in PHENIX^33^. The resulting replisome was used for all downstream model building below.

For model 1, the nucleosome (PDB 3LZ0)^11^ and replisome (above) were rigid body docked into map H. The resulting models were joined and locally adjusted in Coot and then real space refined in PHENIX. For model 2, the nucleosome (PDB 3LZ0) and replisome (above) were rigid body docked into map L. DNA from the nucleosome was removed to match map L, and missing DNA was manually built first in UCSF ChimeraX and then locally adjusted in Coot. The resulting model was real space refined in PHENIX.

For model 3a, the nucleosome (PDB 3LZ0) and replisome (above) were rigid body docked into map Q. DNA from the nucleosome was removed to match map Q, and missing DNA was manually built first in UCSF ChimeraX and then locally adjusted in Coot. The resulting model was real space refined in PHENIX.

For model 3b, model 3a was used as a starting point and the H2A–H2B dimer was removed to match map R and then real space refined in PHENIX.

## Figure generation

Figures were generated using Adobe Illustrator, and UCSF ChimeraX.

## Data availability

The cryo-EM reconstructions and final models will be made available via the Electron Microscopy Data Base and the Protein Data Bank upon completion of peer review.

## Acknowledgements

We thank all members of the Farnung lab, J.C. Walter, and D. Moazed for discussions. We thank S. Shao and D. Sherpa for guidance regarding cryo-EM grid preparation. We thank The Harvard Cryo-EM Center for Structural Biology at Harvard Medical School for support with data collection. We thank S.M. Vos for critical reading and input. F.S. is supported by an NSF Graduate Research Fellowship (DGE 2140743). J.W.M is supported by the Alfred and Joan Goldberg Education and Fellowship Fund for Cell Biology (Harvard Medical School, Department of Cell Biology). L.F. is supported by the NIH Director’s New Investigator Award (DP2-ES036404), the Harvard Medical School Grinnell Fund, and the William F. Milton Award (Harvard University).

## Author contributions

F.S. and J.W.M purified protein components. F.S. conducted all biochemical experiments. F.S. and J.W.M. developed complex formation procedures and F.S. prepared samples for cryo-EM. F.S. processed all cryo-EM data with input from J.W.M. F.S. and J.W.M. prepared figures. L.F. designed and supervised research. F.S. and L.F. wrote the manuscript with input from all authors.

---

**Extended Data Fig. 1.**
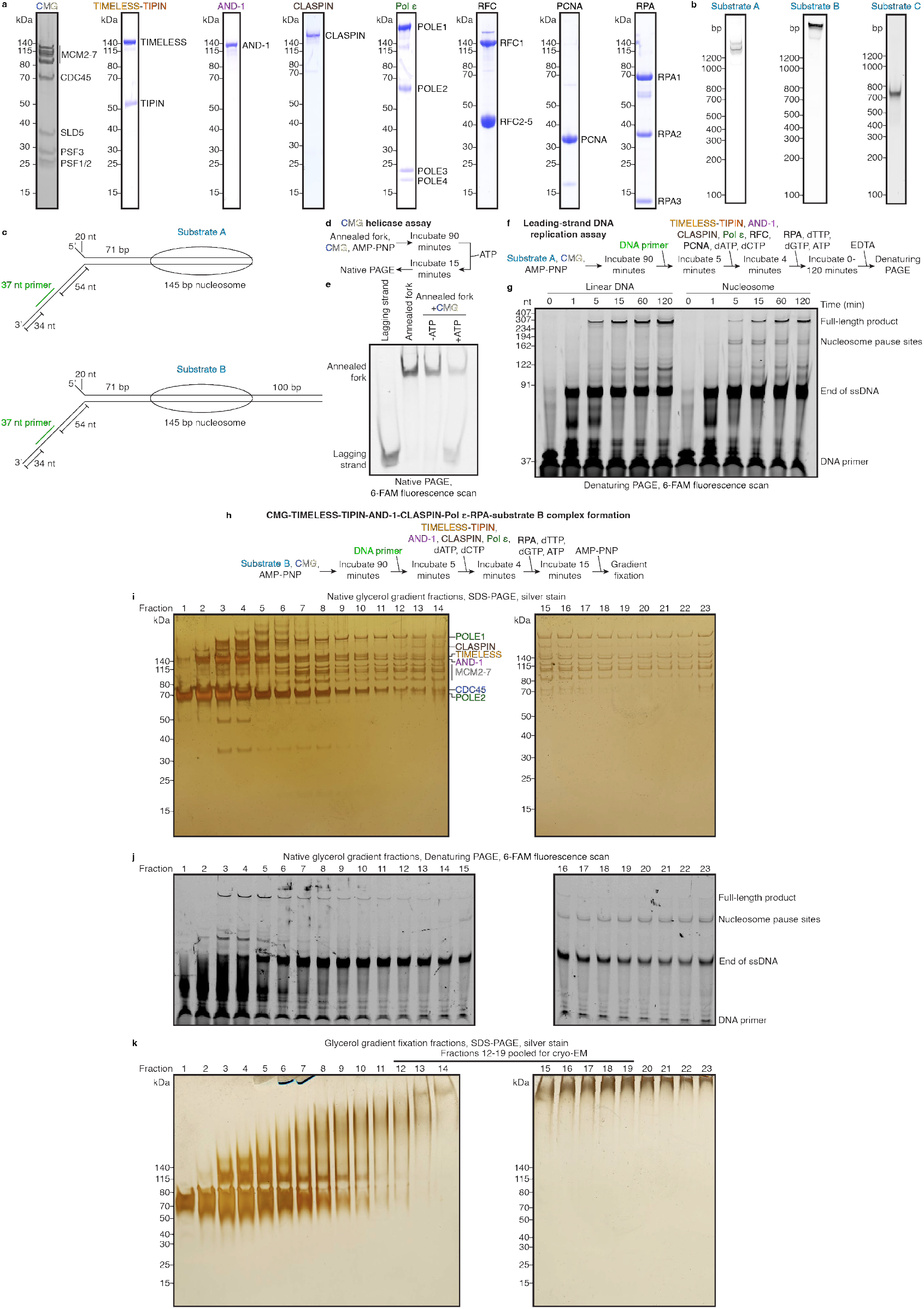
Purified protein components, prepared nucleosomes, helicase and leading-strand DNA replication assays, and CMG-TIME-LESS-TIPIN-AND-1-CLASPIN-Pol ε-RPA-substrate B complex formation. **a**, SDS-PAGE of purified proteins. Molecular weights are indicated. **b**, Native PAGE of nucleosomes used for biochemical and structural studies. **c**, Schematic of nucleosomal substrates A and B, used for leading-strand DNA replication assays and CMG-TIMELESS-TIPIN-AND-1-CLASPIN-Pol ε-RPA-substrate B complex formation, respectively. **d**, Schematic of CMG helicase assay. **e**, Native PAGE of CMG helicase assay. **f**, Schematic of leading-strand DNA replication assay. **g**, Denaturing PAGE of leading-strand DNA replication assay on linear DNA and nucleosomal substrate. **h**, Schematic of CMG-TIMELESS-TIPIN-AND-1-CLASPIN-Pol ε-RPA-substrate B complex formation. **i**, Silver-stained SDS-PAGE of native glycerol gradient fractions. **j**, 6-FAM fluorescence-scanned denaturing PAGE of native glycerol gradient fractions. **k**, Silver-stained SDS-PAGE of glycerol gradient fixation fractions.

**Extended Data Fig. 2.**
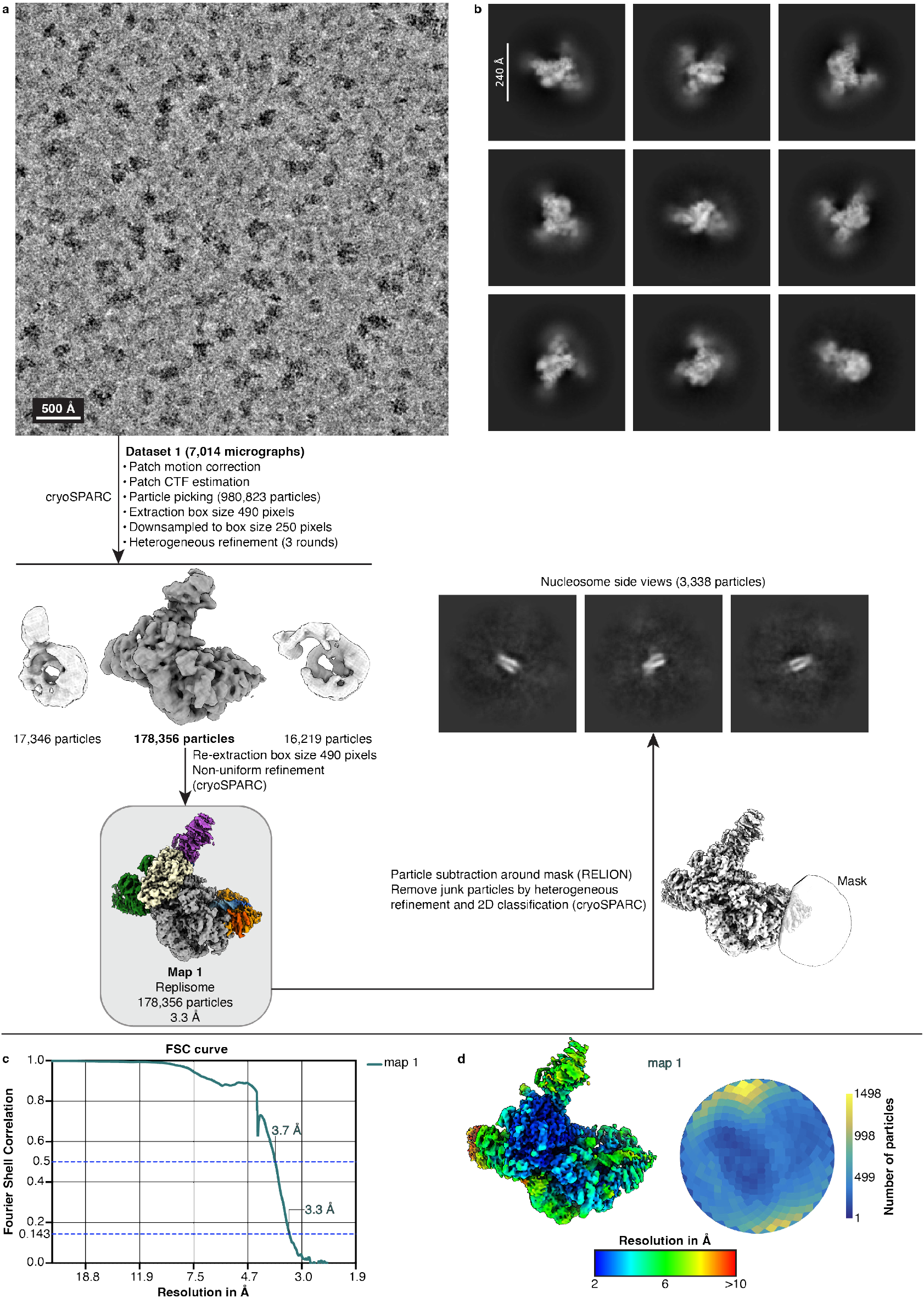
Cryo-EM data analysis, FSC curve, local resolution, and angular distribution plot for CMG-TIMELESS-TIPIN-AND-1-CLASPIN-Pol ε-RPA-substrate B complex. **a**, Representative low-pass filtered micrograph and classification tree. Scale bar for micrograph is 500 Å. **b**, Representative 2D classes of complex. Scale bar is 240 Å. **c**, FSC curve of map 1. **d**, Local resolution of map 1 and corresponding angular distribution plot.

**Extended Data Fig. 3.**
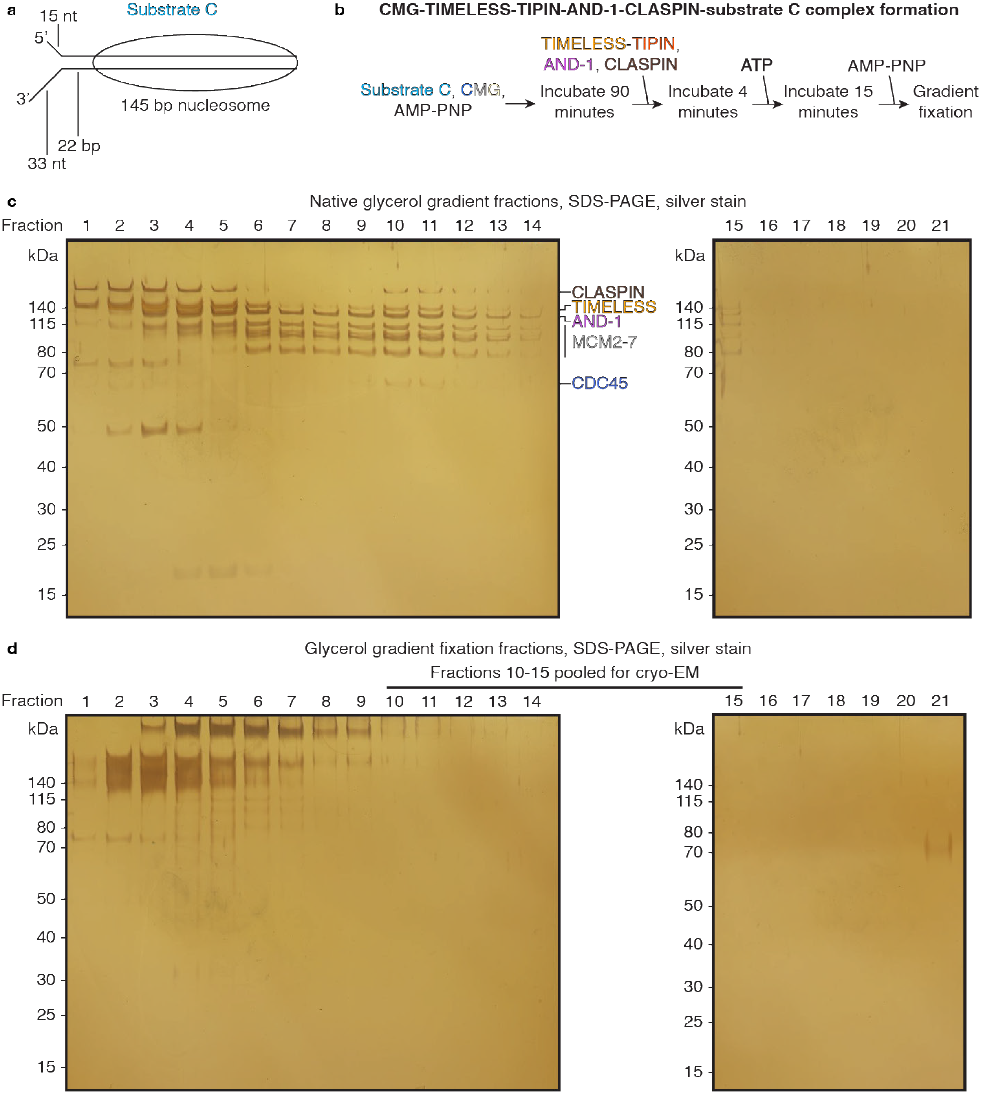
CMG-TIMELESS-TIPIN-AND-1-CLASPIN-substrate C complex formation. **a**, Schematic of nucleosomal substrate C. **b**, Schematic of CMG-TIMELESS-TIPIN-AND-1-CLASPIN-substrate C complex formation. **c**, Silver-stained SDS-PAGE of native glycerol gradient fractions. **d**, Silver-stained SDS-PAGE of glycerol gradient fixation fractions.

**Extended Data Fig. 4.**
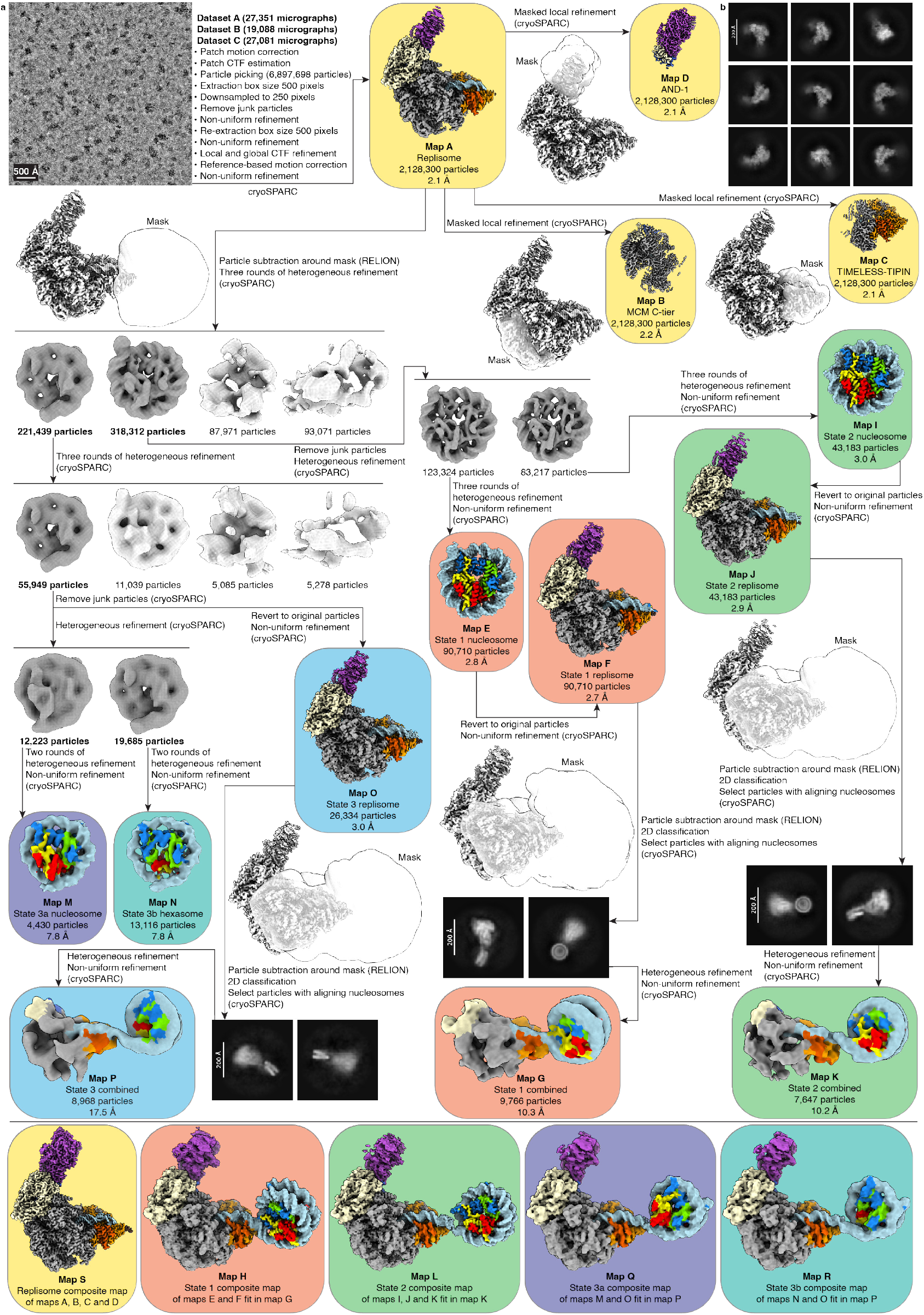
Cryo-EM data analysis for CMG-TIMELESS-TIPIN-AND-1-CLASPIN-substrate C complex. **a**, Representative low-pass filtered micrograph and classification tree. Scale bar for micrograph is 500 Å. Final map background colors indicate which maps were used to construct each final composite map. **b**, Representative 2D classes of complex. Scale bar is 200 Å.

**Extended Data Fig. 5.**
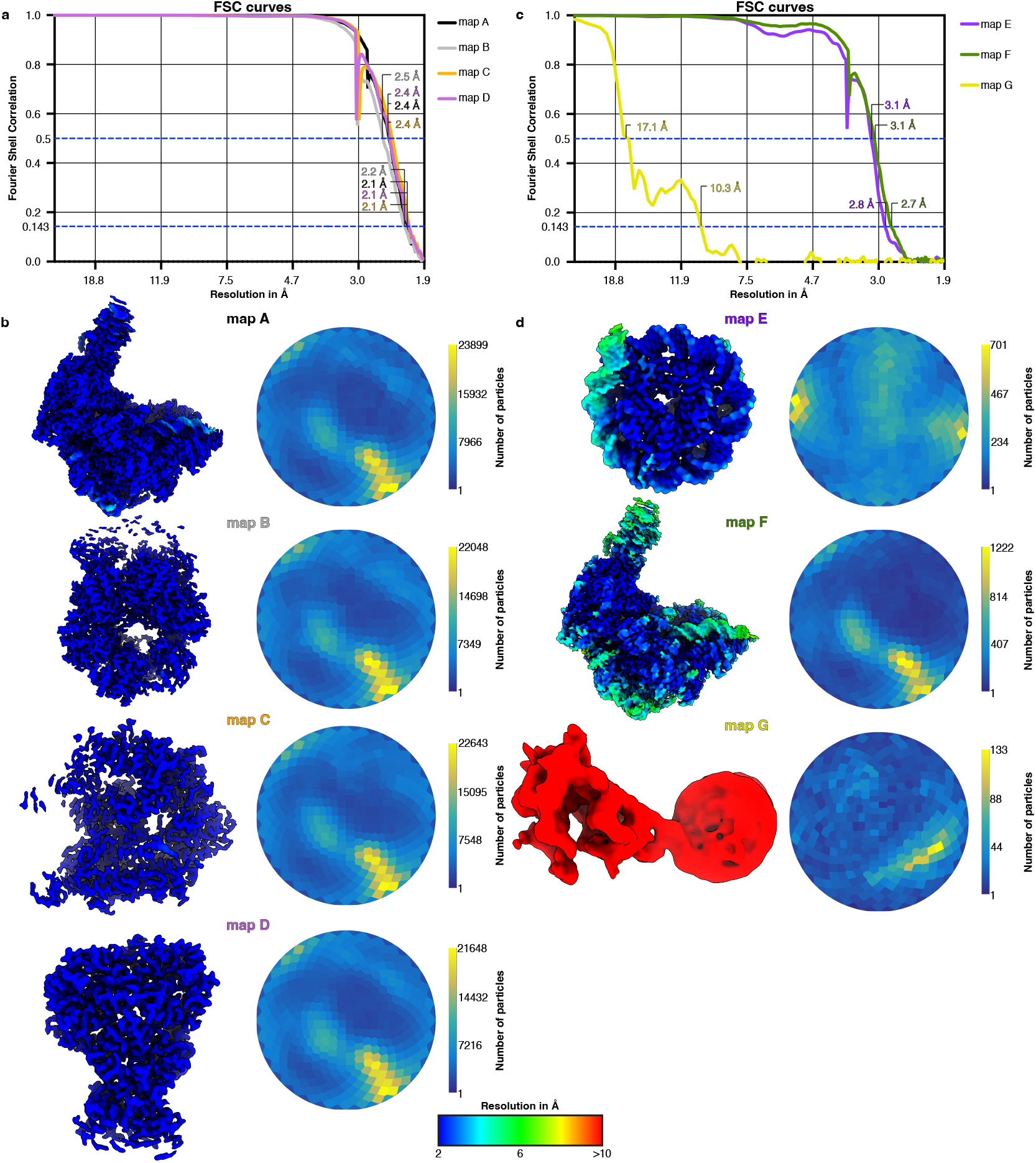
FSC curves, local resolutions, and angular distribution plots for CMG-TIMELESS-TIPIN-AND-1-CLASPIN-substrate C complex. **a**, FSC curves of maps A-D. **b**, Local resolution of maps A-D and corresponding angular distribution plots. **c**, FSC curves of maps E-G. **d**, Local resolution of maps E-G and corresponding angular distribution plots.

**Extended Data Fig. 6.**
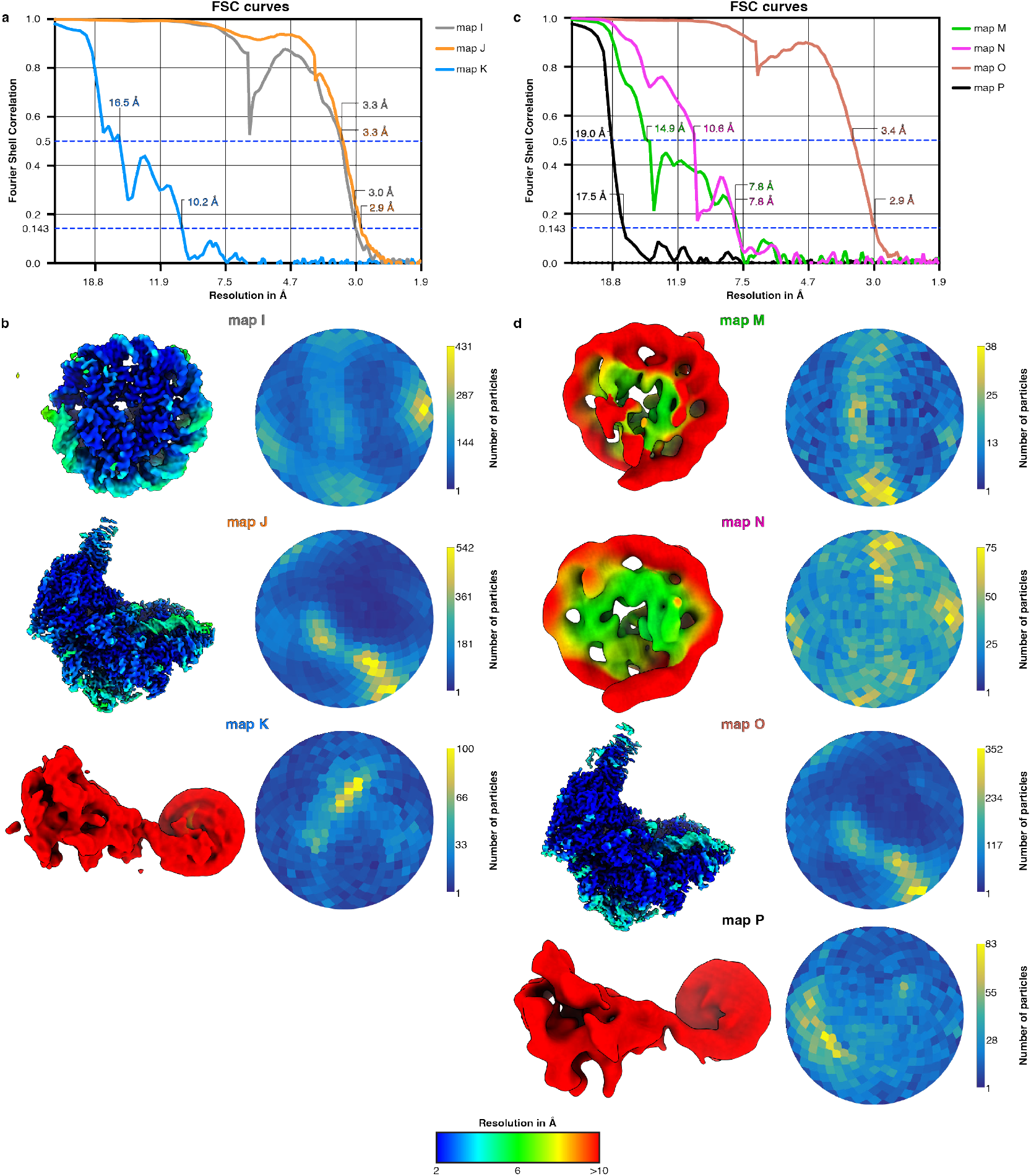
FSC curves, local resolutions, and angular distribution plots for CMG-TIMELESS-TIPIN-AND-1-CLASPIN-substrate C complex, continued. **a**, FSC curves of maps I-K. **b**, Local resolution of maps I-K and corresponding angular distribution plots. **c**, FSC curves of maps M-P. **d**, Local resolution of maps M-P and corresponding angular distribution plots.

**Extended Data Fig. 7.**
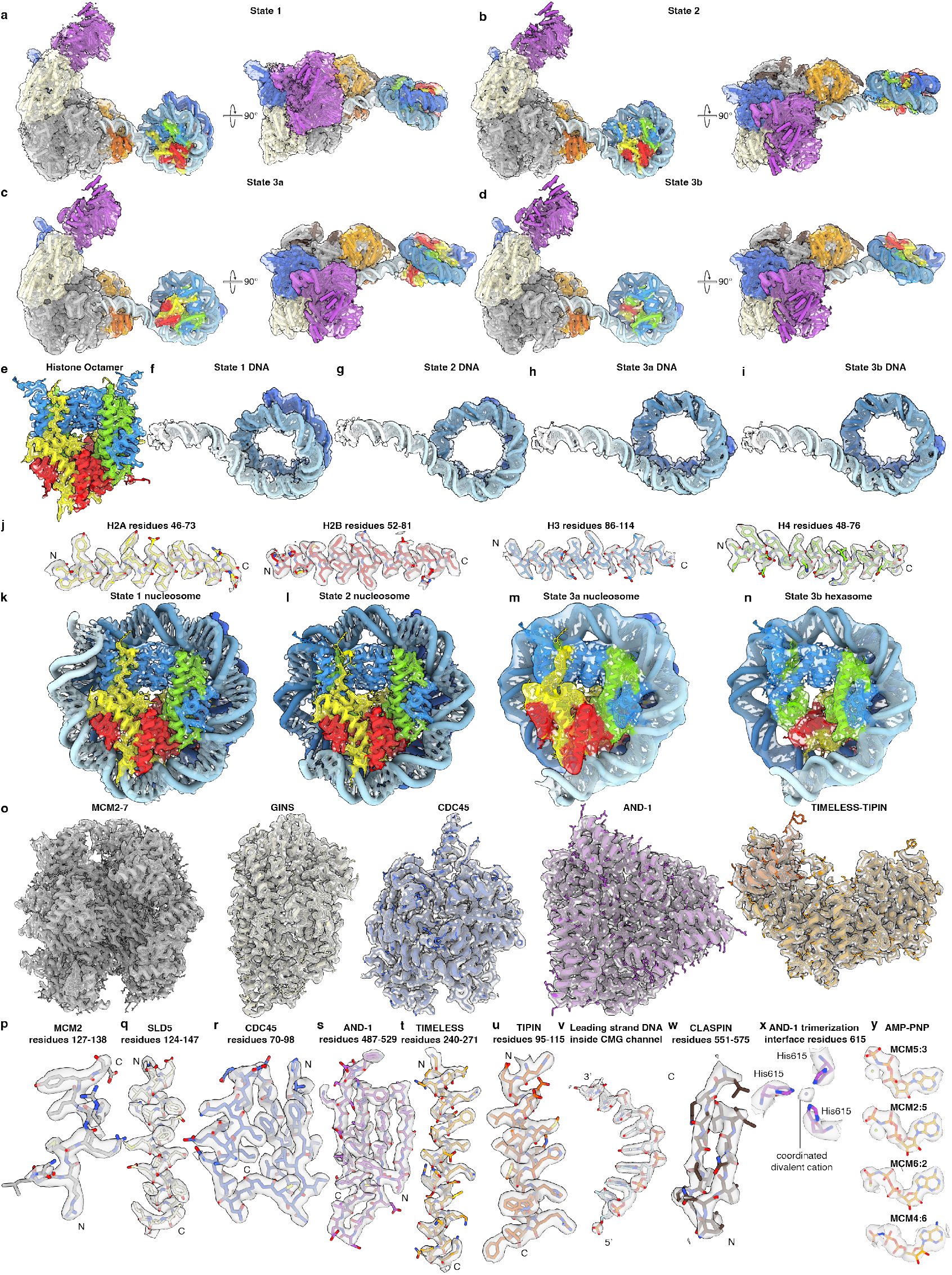
Representative densities. **a**, Coulomb potential map of state 1 replisome-nucleosome complex (map H) with fitted model. **b**, Coulomb potential map of state 2 replisome-nucleosome complex (map L) with fitted model. **c**, Coulomb potential map of state 3a replisome-nucleosome complex (map Q) with fitted model. **d**, Coulomb potential map of state 3b replisome-hexasome complex (map R) with fitted model. **e**, Coulomb potential map of histone octamer (map E) with fitted model. **f**, Coulomb potential map of state 1 DNA (map H) with fitted model. **g**, Coulomb potential map of state 2 DNA (map L) with fitted model. **h**, Coulomb potential map of state 3a DNA (map Q) with fitted model. **i**, Coulomb potential map of state 3b DNA (map R) with fitted model. **j**, Coulomb potential map of histones H2A (residues 46-73), H2B (residues 52-81), H3 (residues 86-114), and H4 (residues 48-76) (map E) with fitted models. **k**, Coulomb potential map of state 1 nucleosome (map E) with fitted model. **l**, Coulomb potential map of state 2 nucleosome (map I) with fitted model. **m**, Coulomb potential map of state 3a nucleosome (map M) with fitted model. **n**, Coulomb potential map of state 3b hexasome (map N) with fitted model. **o**, Coulomb potential map of MCM2-7, GINS, CDC45, AND-1, and TIMELESS-TIPIN (map S) with fitted models. **p**, Coulomb potential map of MCM2 residues 127-138 (map S) with fitted model. **q**, Coulomb potential map of SLD5 residues 124-147 (map S) with fitted model. **r**, Coulomb potential map of CDC45 residues 70-98 (map S) with fitted model. **s**, Coulomb potential map of AND-1 residues 487-529 (map S) with fitted model. **t**, Coulomb potential map of TIMELESS residues 240-271 (map S) with fitted model. **u**, Coulomb potential map of TIPIN residues 95-115 (map S) with fitted model. **v**, Coulomb potential map of leading strand DNA inside CMG channel (map S) with fitted model. **w**, Coulomb potential map of CLASPIN residues 551-575 (map S) with fitted model. **x**, Coulomb potential map of AND-1 trimerization interface residues 615 (map S) with fitted model. **y**, Coulomb potential map of AMP-PNP at MCM5:3, MCM2:5, MCM6:2 and MCM4:6 ATPase sites (map S) with fitted models.

**Extended Data Fig. 8.**
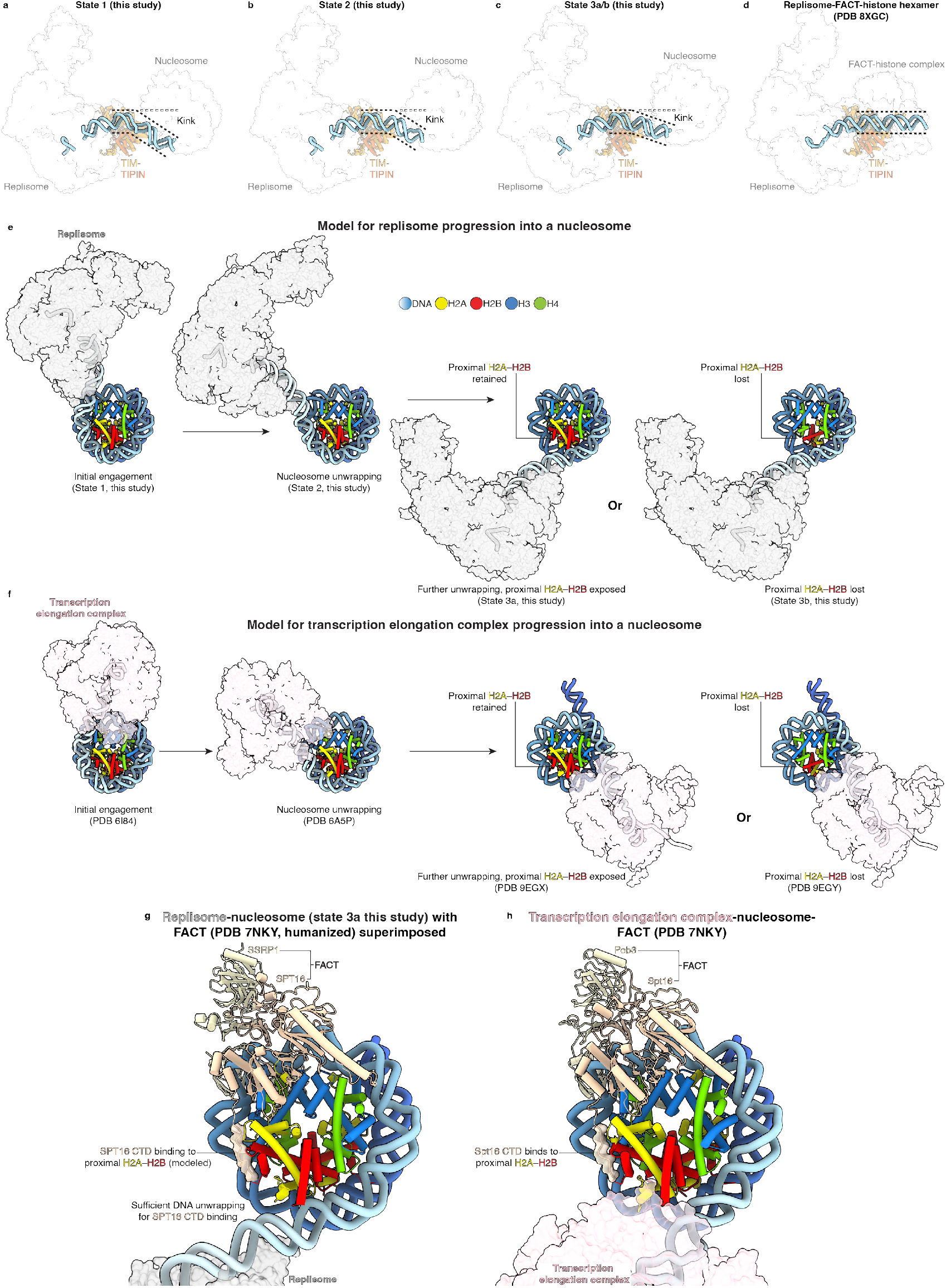
DNA kink at TIMELESS-TIPIN-nucleosome interface, model for replisome progression into a nucleosome and comparison to transcription, and FACT superposition onto replicated nucleosome. **a-d**, Comparison of the downstream DNA trajectory in replisome-nucleosome structures (this study) with the trajectory in a later intermediate of replisomal histone transfer in yeast. **e**, Model for replisome progression into a nucleosome based on the four structures from this study. **f**, Corresponding model for transcription elongation complex progression into a nucleosome based on published structures. **g**, State 3a replisome-unwrapped nucleosome complex with superimposed FACT (PDB 7NKY, humanized). **h**, Structure of a transcription elongation complex engaged with a FACT-bound unwrapped nucleosome (PDB 7NKY) for comparison.

